# Ion modulation restores local synaptic function in APOE4 neuronal networks despite persistent impairments in global network organization

**DOI:** 10.64898/2026.09.08.750089

**Authors:** Marthe Bendiksvoll Grønlie, Anna Mikalsen Kollstrøm, Axel Sandvig, Ioanna Sandvig

## Abstract

The apolipoprotein E ε4 (APOE4) allele is the strongest genetic risk factor for developing sporadic Alzheimer’s disease (AD), which accounts for the vast majority of AD cases. Ion dysfunction has emerged as a central mechanism involved in APOE4-related pathology, reflecting the importance of ion homeostasis for cell membrane excitability, synaptic transmission, and plasticity. We have previously demonstrated that ion dysfunction contributes to early AD-related network vulnerability, and that engaging synaptic plasticity mechanisms via ion modulation can restore network balance in neurodegenerative disease. Here, we investigated whether synaptic mobilization by ion modulation could ameliorate APOE4-associated dysfunction in human neuronal networks compared with isogenic APOE3 controls. APOE4 networks exhibited progressive structural and functional deterioration, manifesting as increased neurite length, reduced neurite branching, and reduced firing rate and number of active recording electrodes. This was accompanied by compensatory hypersynchronization, rendering networks vulnerable to pathological spread, and reduced synaptic AMPAR levels compared with isogenic APOE3 control networks. Modulating ion homeostasis restored the local firing dynamics and stabilized synaptic AMPAR levels. However, the elevated synchronization and aberrant morphology of APOE4 networks were not alleviated by engaging synaptic plasticity mechanisms, demonstrating that APOE4 networks retain a capacity for adaptive plasticity enabling restoration of local neuronal function, but that this recovery does not extend to higher-order network organization.

## Introduction

The apolipoprotein E ε4 (APOE4) allele is the most prevalent genetic risk factor for developing Alzheimer’s disease (AD). In contrast to the rare early-onset familial type of AD, which involves disease-causing mutations in the amyloid precursor protein, presenilin-1, or presenilin-2 gene, more than 95% of cases are sporadic, without fully understood causes involving genetic, environmental, and age-related factors (Pinals and Tsai, 2022). The strongest genetic determinant for sporadic AD is the ε4 isoform of the APOE protein, with disease risk increasing according to the number of ε4 alleles inherited. Compared with homozygous APOE3, which entails a neutral risk for developing AD, carrying two copies of the APOE4 allele increases the risk of AD 8-15 times (Pinals and Tsai, 2022). The APOE protein plays multiple roles in both neurons and glia, regulating lipid homeostasis, energy metabolism, the autophagy-lysosomal function, inflammation, and the development of hallmark amyloid-β (Aβ) and tau pathology through several pathways reviewed previously by Chen et al. (2025b).

Increasing evidence suggests that synaptic impairment and network dysfunction are common features across neurodegenerative diseases, including, AD (Li et al., 2025) (Chen et al., 2025a) (Groenlie et al., 2026), amyotrophic lateral sclerosis (ALS) (Fiskum et al., 2026) (Kollstrøm et al., 2025a) (Verstraete et al., 2010) (Martínez-Silva et al., 2018), and Parkinson’s disease (Valderhaug et al., 2024) (Ko et al., 2018) (Swanson et al., 2023). Ion homeostasis underpins synaptic function and accounts for a substantial proportion of neuronal energy consumption through the continuous active transport required to maintain ionic gradients across the cell membrane (Bhoi et al., 2025). Disruption of ion homeostasis impairs membrane excitability, neurotransmission, and synaptic plasticity, leading to increased metabolic demand, calcium toxicity, oxidative stress, inflammation, and disruption of intracellular signalling cascades (Bhoi et al., 2025) (Li et al., 2024), all of which contribute to neurodegenerative disease (Zhang et al., 2024).

Synaptic ion channel dysfunction has been established as a key contributor to early AD-related network vulnerability (Bhoi et al., 2025) (Adeoye and Ullah, 2025). In a previous study, we identified dysregulation of monoatomic inorganic cation transport together with impaired synaptic transmission and markedly reduced firing rates in human APOE4 neuronal networks (Groenlie et al., 2026). These findings suggest that disrupted ion homeostasis contributes to network hypoexcitability and that restoring neuronal excitability may represent a viable strategy to normalize network activity in AD. This is further supported by our previous work in ALS, in which synaptic potentiation by potassium channel blockage restored network activity despite distinct disease-specific pathology (Kollstrøm et al., 2025b) (Kollstrøm et al., 2025a). Given these previous findings, and the central role of potassium channels in regulating neuronal excitability and synaptic transmission, we hypothesized that potassium channel blockage in AD networks could improve synaptic function and thereby restore the functional deficits observed in APOE4 neuronal networks (Aniksztejn and Ben-Ari, 1991b) (Huang and Malenka, 1993) (Huber et al., 1995) (Aniksztejn and Ben-Ari, 1991a) (Paulsen et al., 1990).

## Methods

### Differentiation of human iPSCs into cortical neurons

Human induced pluripotent stem cells (iPSCs) were obtained from The Jackson Laboratory. To model genetic susceptibility to AD, the homozygous APOE4 cell line JIPSC001142 (Safiri et al., 2024) was used and compared with its parental isogenic control line KOLF2.1J (JIPSC001000; passage 4; male donor aged 55–59 years) harbouring two APOE3 alleles (Pantazis et al., 2022). APOE3 is associated with a neutral risk for developing AD (Safiri et al., 2024). iPSCs were expanded and differentiated into cortical neurons following a protocol by Dannert et al. (2023), with adaptations previously described in Groenlie et al. (2026). Cells were plated onto different plate formats depending on downstream applications: 8-well chamber slides (Nunc LabTek, 177445) for immunocytochemistry, Cytoview 12-well microelectrode arrays (MEAs) (Axion Biosystems, M768-tMEA-12W) for electrophysiological recordings, and standard 6-well plates (Falcon, Corning, 353046) for western blot. This day of the final plating was considered 0 days in vitro (DIV). Neurons were maintained by replacing 50% of the cell medium every two days as described in Dannert et al. (2023).

### Engaging synaptic plasticity by ion modulation

Based on our previous findings that ion imbalance and synaptic dysfunction contribute to network vulnerability in AD (Groenlie et al., 2026), engagement of synaptic plasticity mechanisms by ion modulation was performed in half of the networks from each group by applying a potassium-channel blocker, tetraethylammonium (TEA), at concentrations and durations previously described in Kollstrøm et al. (2025b). Briefly, before three subsequent full media changes, 25 mM TEA was applied for 10 min. TEA causes prolonged action potential duration and repolarization phase, transiently enhanced glutamate release, and increased postsynaptic calcium influx (Aniksztejn and Ben-Ari, 1991b) (Huang and Malenka, 1993) (Huber et al., 1995) (Aniksztejn and Ben-Ari, 1991a) (Paulsen et al., 1990). Due to these effects, TEA has also been used to induce long-term potentiation (LTP) (Aniksztejn and Ben-Ari, 1991b) (Huang and Malenka, 1993) (Huber et al., 1995) (Kollstrøm et al., 2025b). Based on our previous study demonstrating an APOE4-dependent shift in network activity from 52 DIV (Groenlie et al., 2026), TEA was applied at 47, 49, and 51 DIV to prevent or delay the previously observed impairments in network function. Networks used for immunocytochemistry and western blot were fixed or harvested one day following the final ion modulation to obtain parallel results with the ongoing schedule for longitudinal electrophysiological recordings. The full experimental timeline is illustrated in Fig. 1.

**Fig. 1.**
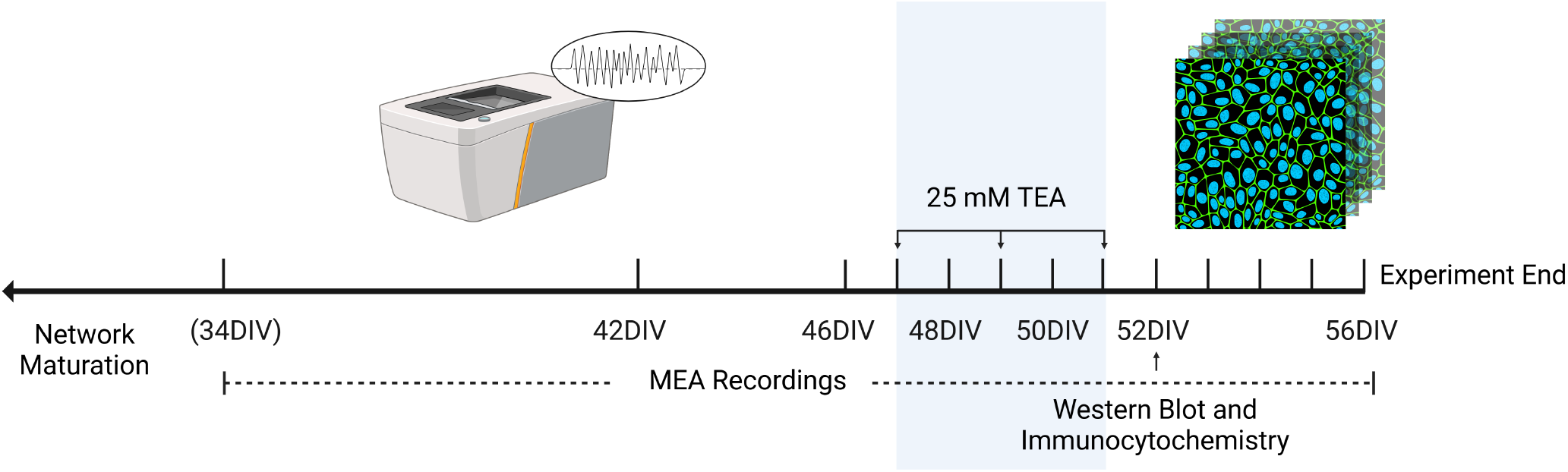
Experimental timeline. Human induced pluripotent stem cell-derived neuronal networks were established on microelectrode arrays (MEAs), chamber slides, and 6-well plates for longitudinal electrophysiological recordings, immunocytochemistry and western blot, respectively. At 47, 49, and 51 days in vitro (DIV) (shaded area), ion modulation was performed in half of the networks using 25 mM tetraethylammonium (TEA). After test recordings at 34 DIV, regular recordings were obtained every four days from 42-56 DIV, except during the perturbation when recordings were obtained every two days. At 52 DIV, networks were fixed and harvested for immunocytochemistry and western blot. Figure created using BioRender.

### Electrophysiological recordings and analysis

Longitudinal neuronal network activity was monitored by 20 min long extracellular MEA recordings using the Axion Maestro Pro MEA system (Axion Biosystems, GA, USA). Data acquisition was carried out with AxIS Navigator (version 3.10.3.6) at a sampling frequency of 12.5 kHz. Recordings were obtained from n=12 wells per condition. Prior to each recording, networks were left inside the Maestro Pro for 10 min to stabilize activity. Recordings were collected between 34 and 56 DIV. Baseline test recordings were obtained at 34 DIV, before regular recordings were obtained every four days (every two days during TEA application) from 42 to 56 DIV. Recordings were excluded from the analysis if they contained fewer than seven spikes per minute or fewer than five active electrodes. Spike and burst detection was performed using the AxIS Navigator analysis tool with parameter settings as previously described in Groenlie et al. (2026). Due to a low number of active electrodes in some groups, the coherence index was used to assess network synchrony, calculated as the ratio between the standard deviation and the mean of the instantaneous firing rate (Wang, 2002). A high coherence index indicates higher network synchronization. Further processing of spike and burst statistics was performed in Matlab (version R2023a) (MathWorks, 2023). Outliers were removed if located more than three inter-quartile ranges outside the middle 50% of the data without any clear pattern in belonging to group or time point. The resulting number of networks included in the analysis per group and time point are listed in Supplementary Table S3.

### Statistical analysis of electrophysiology data

Statistical analysis of electrophysiological data was performed using IBM SPSS Statistics (version 30.0.0.0 (172)) (IBM Corp, 2025). Generalized linear mixed models were fitted to determine whether the trajectories of neuronal activity differed between groups. To model trajectories before and after ion modulation, all time points except those during the perturbation phase at 48 and 50 DIV were included in the analysis. The distribution of each electrophysiological metric was evaluated for each group prior to modelling. Individual networks were treated as subjects, time points were included as repeated measures, and a first-order autoregressive covariance structure was specified to account for temporal dependency among repeated observations within wells. The different electrophysiology metrics served as dependent variables. Fixed effects included group, time point, and the interaction between group and time point, together with model intercepts. Effective degrees of freedom were estimated using the Satterthwaite approximation. Differences between groups were evaluated using pairwise contrasts while holding time points constant at their mean values. To control for multiple comparisons, p-values were adjusted using sequential Bonferroni correction. The threshold for statistical significance was set to adjusted p = .05. Model performance was evaluated using the Akaike and Bayesian information criteria. The output data generated in Matlab and SPSS were imported into RStudio (RStudio IDE 2025.05.0+496) (R Core Team, 2021) for processing and visualisation using the packages haven (version 2.5.5), readxl (version 1.4.5), dplyr (version 1.1.4), tidyr (version 1.3.1), and ggplot2 (version 3.5.2) (Wickham and Bryan, 2025) (Wickham et al., 2025a) (Wickham et al., 2025c) (Wickham et al., 2025b) (Wickham, 2016).

### Synaptosome isolation and western blot

Synaptic α-amino-3-hydroxy-5-methyl-4-isoxazolepropionic acid receptor (AMPAR) levels were assessed by western blot, given that potassium channel blockage can induce LTP (Aniksztejn and Ben-Ari, 1991b) (Huang and Malenka, 1993) (Huber et al., 1995) (Kollstrøm et al., 2025b) (Caya-Bissonnette and Béïque, 2024). Synaptosomes were isolated from the networks at 52 DIV as previously described in Groenlie et al. (2026), and western blot was performed by the Proteomics and Modomics Experimental Core (PROMEC) facility at the Norwegian University of Science and Technology (NTNU). Samples were prepared in LDS loading buffer and 1D PAGE was performed in 10% NuPage Novex Bis-Tris gels using MOPS run buffer. Gels were electroblotted on nitrocellulose using turboblot (BioRad). Membranes were blocked for 1h in PBS, 0.1% Tween (PBST), 5% fat-free dry milk and incubated in primary antibody GluR1 (Thermo Fisher, MA5-27694) for 1h in blocking buffer. Tuj1 (EP1569Y, Abcam, ab52623) was used as loading control. After 3*10 min washing in PBST, membranes were further incubated for 1h in secondary IRDYE-conjugated antibodies (700CW or 800CW) in PBST, washed 3*10 min in PBST, 1*10 min PBS and visualized using an Odyssey Imaging system (Licor). Because we compared more than two independent groups with non-normal distributions, the Kruskal-Wallis H test was used for statistical comparisons.

### Immunocytochemistry

Neuronal networks on 8-well chamber slides were fixed at 52 DIV according to the protocol by Richter et al. (2018), and subsequently immunolabeled as previously described in Groenlie et al. (2026) using the primary and secondary antibodies listed in Table 1. After secondary antibody incubation, cell nuclei were counterstained using 8 *µ*M Hoechst (Thermo Fisher, 33342) for 10 min. Fluorescence imaging was performed using an EVOS M7000 microscope (Invitrogen) with a 20x/0,45 NA EVOS AMEP 4982 objective and LED light cubes DAPI (AMEP4650), GFP (AMEP4651), TX-Red (AMEP4655), and CY5 (AMEP4656). Image processing was performed in Fiji/ImageJ (v1.54r).

**Table 1.** Primary and Secondary Antibodies Used.

| Target | Primary antibody (concentration, product) | Secondary antibody (product). All at 1:500 concentration |
| --- | --- | --- |
| Axons, neurofilament | Chicken anti-NfH (1:1000, Abcam, ab4680) | Goat anti-chicken Alexa Fluor 488 (Invitrogen, A11039) |
| Dendrites, microtubules | Chicken anti-MAP2 (1:2000, Abcam, ab5392) | Goat anti-chicken Alexa Fluor 488 (Invitrogen, A11039) |
| Mid-layer cortical neurons | Mouse anti-SATB1+2 (1:100, Abcam, ab51502) | Goat anti-mouse Alexa Fluor 488 (Invitrogen, A28175) |
| GABA synthesis | Mouse anti-GAD67 (1:100, Sigma-Aldrich, MAB5406) | Goat anti-mouse Alexa Fluor 488 (Invitrogen, A28175) |
| Amyloid-β peptides | Mouse anti-McSA1 (1:750, Medimabs, MM-0015-P) | Goat anti-mouse Alexa Fluor 647 (Abcam, ab150115) |
| P-tau (Ser202, Thr205) | Mouse anti-AT8 (1:500, Thermo Fisher, MN1020) | Goat anti-mouse Alexa Fluor 647 (Abcam, ab150115) |
| NMDA receptors | Mouse-GluN1 (1:100, Sigma-Aldrich, SAB5200546) | Goat anti-mouse Alexa Fluor 647 (Abcam, ab150115) |
| AMPA receptors | Mouse-GluR1 (1:100, Thermo Fisher, MA5-27694) | Goat anti-mouse Alexa Fluor 647 (Abcam, ab150115) |
| Glutamate packaging | Rabbit anti-vGluT1 (1:100, Abcam, ab227805) | Goat anti-rabbit Alexa Fluor 568 (Invitrogen, A11011) |
| Astrocytes | Rabbit anti-GFAP (1:500, Agilent, Z0334) | Goat anti-rabbit Alexa Fluor 568 (Invitrogen, A11011) |
| Deep-layer cortical neurons | Rat anti-Ctip2 (1:300, Abcam, ab18465) | Goat anti-rat Alexa Fluor 568 (Invitrogen, A11077) |

### Structural quantification

Neurite length, density, and branching were quantified using the Fiji/ImageJ (v1.54p) macro set NeuroConnectivity (Verstraelen et al., 2020) (Verstraelen et al., 2024) (Verschuuren et al., 2019), based on the protocol described in Kollstrøm et al. (2025a). Neuronal networks immunolabeled with MAP2 and Hoechst were imaged using an automated scan protocol with autofocus at each field, imaging 75% (APOE4) or 15% (APOE3) of each well area, avoiding edges. Differences in network morphology and image quality required the acquisition of a greater number of images from the APOE4 groups to obtain comparable sample sizes following visual quality assessment and exclusion. Images were obtained from four wells per group, and pixel sizes were 0.309 *×* 0.309 *µ*m. Before processing, images were manually assessed to remove images with artefacts, i.e. images that were out of focus, contained large debris or edges, and images containing only large clusters and no extending neurites. The resulting number of images per group were as follows: APOE3, n=211; APOE3 TEA, n=218; APOE4, n=250; APOE4 TEA, n=197. Neurite detection was based on (Pani et al., 2014), and nuclei were detected using triangle thresholding. The groups were assessed for normality using Shapiro-Wilk’s test. Statistical comparisons were subsequently performed using Kruskal-Wallis test due to non-normal data, followed by Conover’s test with Bonferroni correction for multiple comparisons. All tests were performed using the Python programming language (Python Software Foundation) with packages Scipy (Virtanen et al., 2020), Pingouin (Vallat, 2018) and Scikit-posthocs (Terpilowski, 2019), and figures generated with Matplotlib and Seaborn.

## Results

### Mature cortical neurons and astrocytes in APOE3 and APOE4 networks

Immunocytochemical characterization at 52 DIV demonstrated successful differentiation of both the APOE3 and the APOE4 iPSCs into mature excitatory and inhibitory cortical neurons and astrocytes (Fig. 2A-H). Expression of VGLUT1 and GAD67 confirmed presence of excitatory and inhibitory neurons, respectively (Fig. 2A-B). Expression of CTIP2 and SATB1+2 indicated deep- and mid-layer cortical neuronal identities, respectively (Fig. 2C-D). Expression of cytoskeletal markers NfH and MAP2, and astrocyte marker GFAP, indicated presence of mature neurons together with astrocytes (Fig. 2A-H). McSA1- and AT8-positive cells were identified in both APOE3 and APOE4 networks (Fig. 2E-H), indicating presence of Aβ peptides and phosphorylated tau (Ser202/Thr205), respectively.

**Fig. 2.**
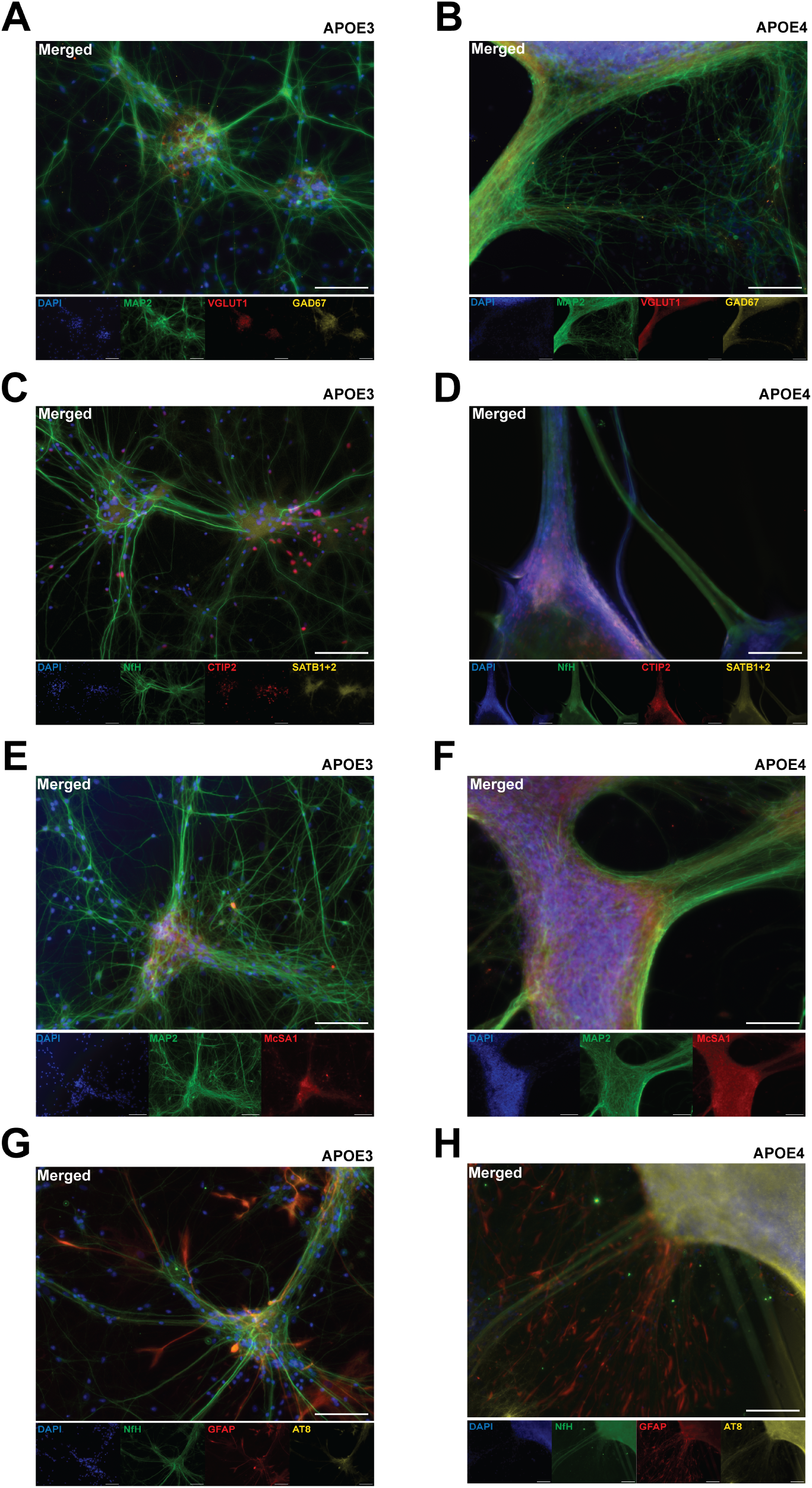
Immunocytochemistry characterizing cell identities in APOE3 and APOE4 networks at 52 days in vitro. (**A-H**) Mature neurons were confirmed by the expression of MAP2 and NfH (**A-B**) Excitatory and inhibitory neuronal identities were confirmed by the expression of VGLUT1 and GAD67, respectively. (**C-D**) Cortical neurons were confirmed by the expression of CTIP2 and SATB1+2. (**E-H**) Aβ peptides and hyperphosphorylated tau were confirmed by the expression of McSA1 and AT8, respectively. (**H**) Astrocytes were confirmed by the expression of GFAP. Scale bar: 100 *µ*m.

### Aberrant clustering and impaired dendritic branching in APOE4 networks

In terms of network morphology, APOE3 neurons organized into small clusters of somata as indicated by stained nuclei, connected by slender and widely dispersed axonal and dendritic projections (Fig. 2A, C, E, G). APOE4 neurons organized into larger and more compact clusters containing both cell nuclei and neurites, interconnected by few, but very thick bundles of axons and dendrites (Fig. 2B, D, F, H). Quantitative assessments of neurite growth at 52 DIV confirmed these observations (Fig. 3), however, the excessive clustering of APOE4 networks introduced some variability in the detection. Accordingly, although the analysis indicated reduced total neurite length (p<.001) (Fig. 3A) and branching (p<.001) (Fig. 3C) along with increased average neurite length (p<.001) (Fig. 3B) in APOE4 compared with APOE3 networks, the extent of these structural differences should be interpreted with some caution. Until 27 DIV, the morphologies of APOE3 and APOE4 networks were comparable, exhibiting evenly dispersed neurites (Supplementary Fig. S1). However, by 41 DIV, light microscopy revealed a pronounced increase in clustering within APOE4 networks, indicating that excessive clustering in APOE4 networks emerged between 27 and 41 DIV.

**Fig. 3.**
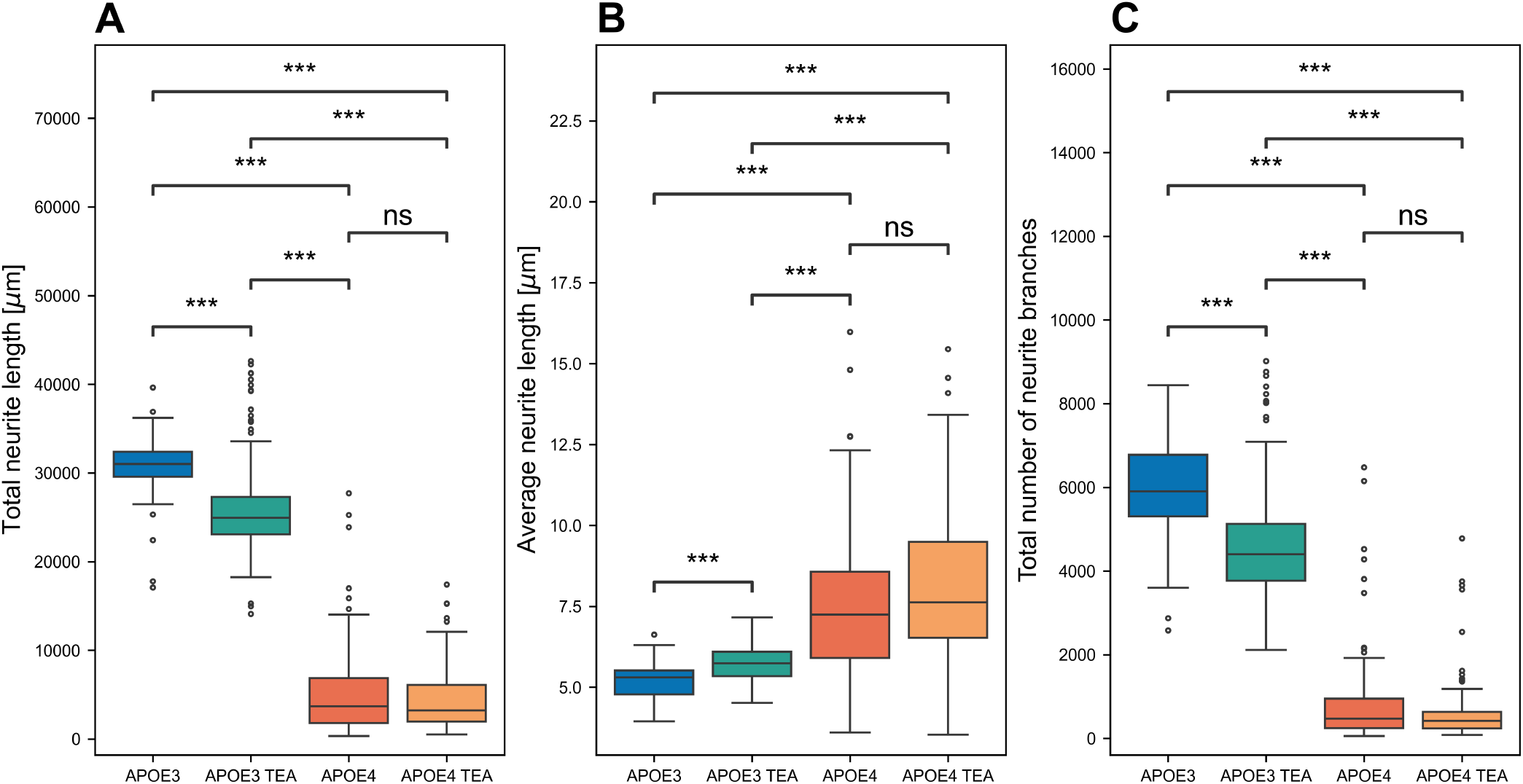
Structural changes in ion-modulated (tetraethylammonium; TEA) and untreated control APOE3 and APOE4 networks at 52 days in vitro (APOE3, n=211; APOE3 TEA, n=218; APOE4, n=250; APOE4 TEA, n=197). (**A-C**) Box plots showing group differences in total neurite length (A), average neurite length (B), and total number of neurite branches (C). Kruskal-Wallis test, followed by Conover’s test for post-hoc comparisons with Bonferroni correction. ***, p<.001; ns, non-significant.

### Structural reorganization of APOE3 networks in response to ion modulation

Modulation of ion dynamics in APOE3 networks significantly reduced branching (p<.001) and total neurite length (p<.001) and increased the average neurite length (p<.001), compared with untreated APOE3 networks (Fig. 3). No significant changes in neurite growth could be identified in APOE4 networks following ion modulation.

### Similar synaptic AMPAR levels in APOE4 networks at baseline and in response to ion modulation

Synaptic AMPAR levels were quantified by western blot at 52 DIV, i.e. one day after the final ion modulation. Both APOE3 and APOE4 networks expressed GluN1 and GluR1 (Fig. 4A-D), confirming the presence of N-methyl-D-aspartate receptors (NMDARs) and AMPARs, respectively. Compared with untreated APOE3 control networks, untreated APOE4 networks exhibited a 1.480-fold reduction in total synaptic AMPAR levels (Fig. 4E). Following ion modulation of APOE3 networks, we found a 3.024-fold decrease in total synaptic AMPAR. Following ion modulation of APOE4 networks, AMPAR levels remained comparable to those of untreated APOE4 networks (Δ=.003). Kruskal-Wallis H test showed no significant differences between any of the groups (χ2=4.924(3), p=.177).

**Fig. 4.**
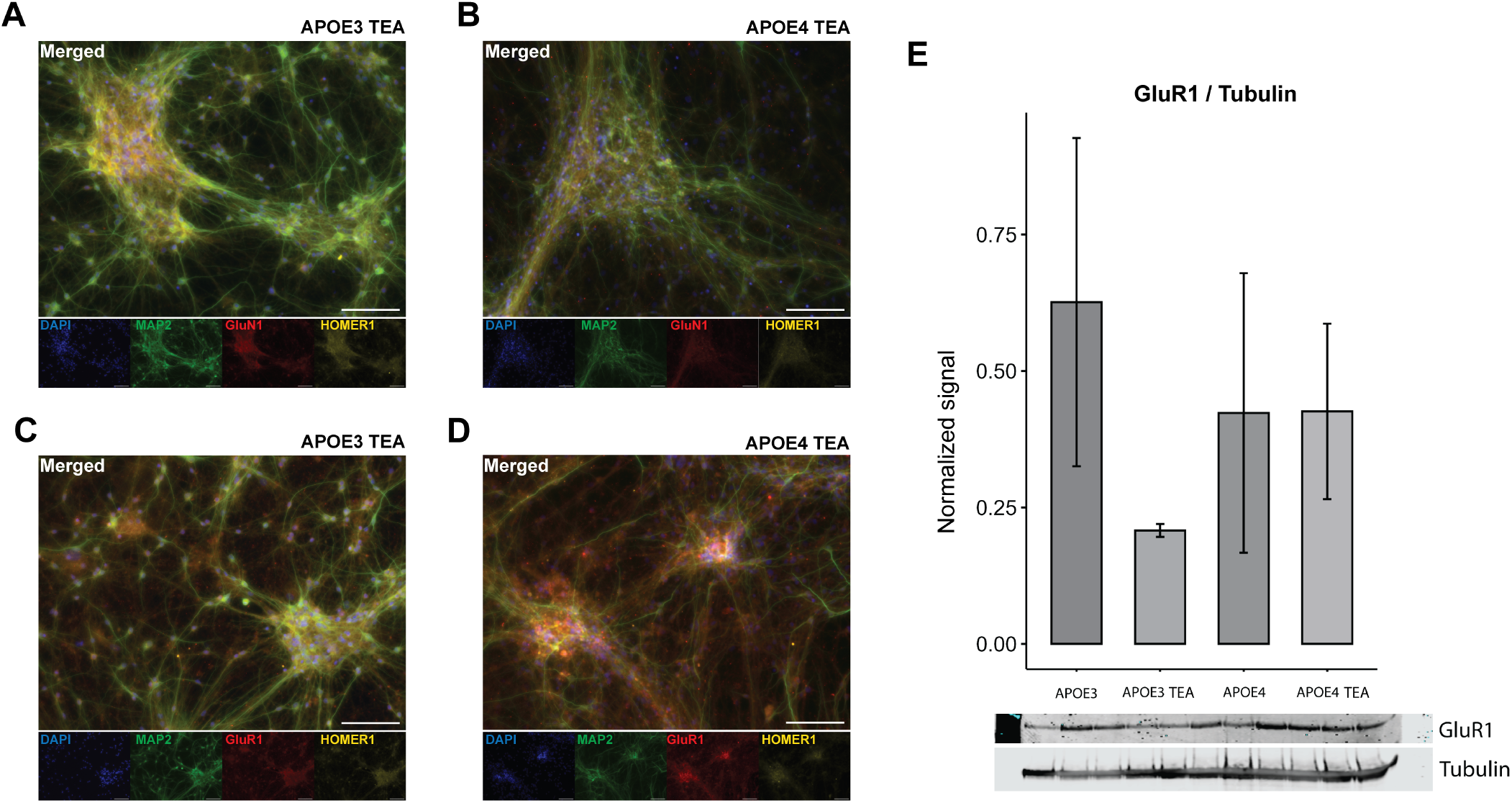
Left panel: Immunocytochemistry of ion-modulated (tetraethylammonium; TEA) APOE3 and APOE4 networks at 52 days in vitro. (**A-D**) NMDA and AMPA receptors were confirmed in both groups by the expression of GluR1 and GluN1, respectively. Scale bar: 100 *µ*m. Right panel: (**E**) Western blot analysis of GluR1, normalized by tubulin (Tuj1). Median signal for APOE3 = .626 (IQR=.301), APOE3 TEA = .208 (IQR=.012), APOE4 = .423 (IQR=.256), and APOE4 TEA = .426 (IQR=.161). Kruskal-Wallis H test showed no significant differences between the groups *χ*2=4.924(3), p=.177

### Partial restoration of APOE4 hypoactivity following ion modulation

Longitudinal MEA recordings of neuronal network activity showed that ion-modulated networks from both groups had a higher median spike amplitude than their untreated counterparts on the day following the final treatment, i.e. at 52 DIV (Fig. 5A-B). While the effect of remained stable in APOE4 networks throughout the observation period, it was no longer evident in APOE3 networks at 56 DIV. At 56 DIV, the amplitude of TEA-treated APOE3 networks had returned to the same level as untreated APOE3 networks. Untreated APOE4 networks consistently showed a higher spike amplitude than untreated APOE3 networks at all measured time points (p=.224). When considering the amplitude trajectories across the entire study period, none of the groups significantly differed. Supplementary Tables S1 and S2 show the fitted generalized linear mixed models for all electrophysiology metrics and groups, with their estimated means, standard errors, confidence intervals, p-values, distributions, and link functions.

**Fig. 5.**
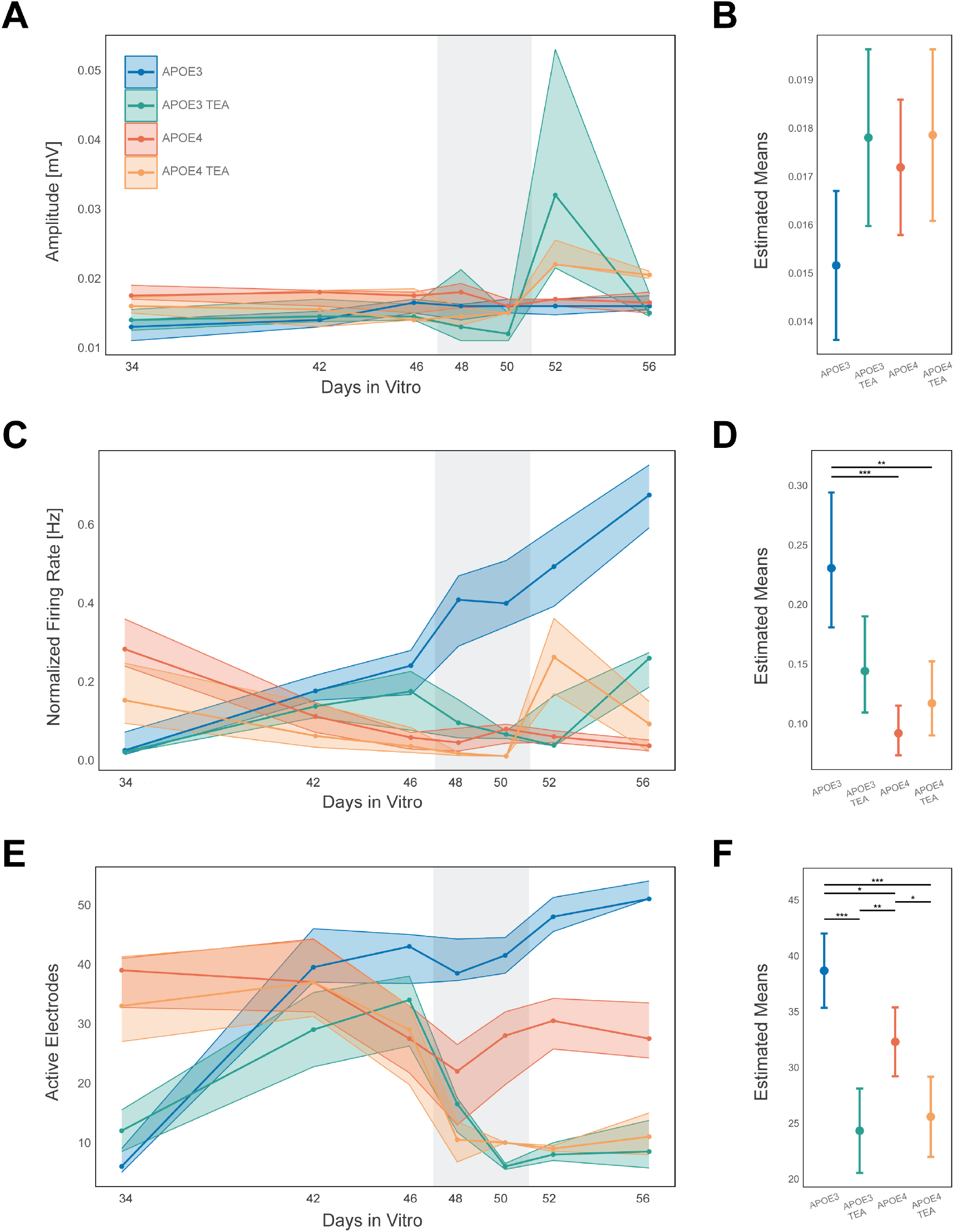
Network activity in ion-modulated (tetraethylammonium; TEA) and untreated control APOE3 and APOE4 networks. (**A-B**) Spike amplitude in mV, (**C-D**) Electrode firing rate in Hz, normalized to the number of active electrodes, and (**E-F**) Number of active electrodes. Left panel: Neuronal network activity over time. Solid lines and circles illustrate median value for all networks in each group. Coloured shaded areas show upper and lower quartile. Vertical shaded gray areas indicate time points of treatment (47, 49, and 51 days in vitro). Group legend is shared between figures. Right panel: Generalized linear mixed models of estimated means with whiskers showing 95% confidence interval for each electrophysiology metrics. See Supplementary Table S3 for n per condition and time point. ***p<.001, **p<01, *p<.05

The median firing rate, normalized to number of active electrodes, increased progressively in untreated APOE3 networks (Fig. 5C-D), whereas the opposite trend was observed in untreated APOE4 controls (p<.001). Ion modulation transiently reduced the firing rate of APOE3 networks during the perturbation and on the following day, after which firing rates increased by the final recording at 56 DIV. Similarly, the firing rate of APOE4 networks decreased slightly during ion modulation, but exhibited a transient peak on the day following the final perturbation.

The number of active electrodes increased progressively over time in untreated APOE3 networks, reaching approximately 50 electrodes by the final recording (Fig. 5E-F). In contrast, in the untreated APOE4 networks, the number of active electrodes fluctuated over time and remained between 25-30 during the final two recordings (p<.05). Ion modulation significantly decreased the number of active electrodes in both groups to comparable levels (p=.632), and this effect persisted until the final recording at 56 DIV. Ion-modulated APOE3 networks had a significantly smaller number of active electrodes than untreated APOE3 networks (p<.001), and ion-modulated APOE4 networks showed a significantly smaller number than untreated APOE4 networks (p<.05).

### Enhanced network synchrony and burst organization following ion modulation

In both groups, synaptic engagement by ion modulation increased the fraction of spikes that were contained in electrode bursts at both time points following TEA perturbation (Fig. 6A-B). When considering the longitudinal slopes of the groups, the effect was only significant for APOE4 networks (p<.001), partly due to the greater variability in APOE3 networks. In untreated controls, APOE4 networks showed a non-significant trend towards a larger fraction of spikes in bursts than APOE3 networks (p=.132). Ion modulation further increased this difference between the groups (p<.01).

**Fig. 6.**
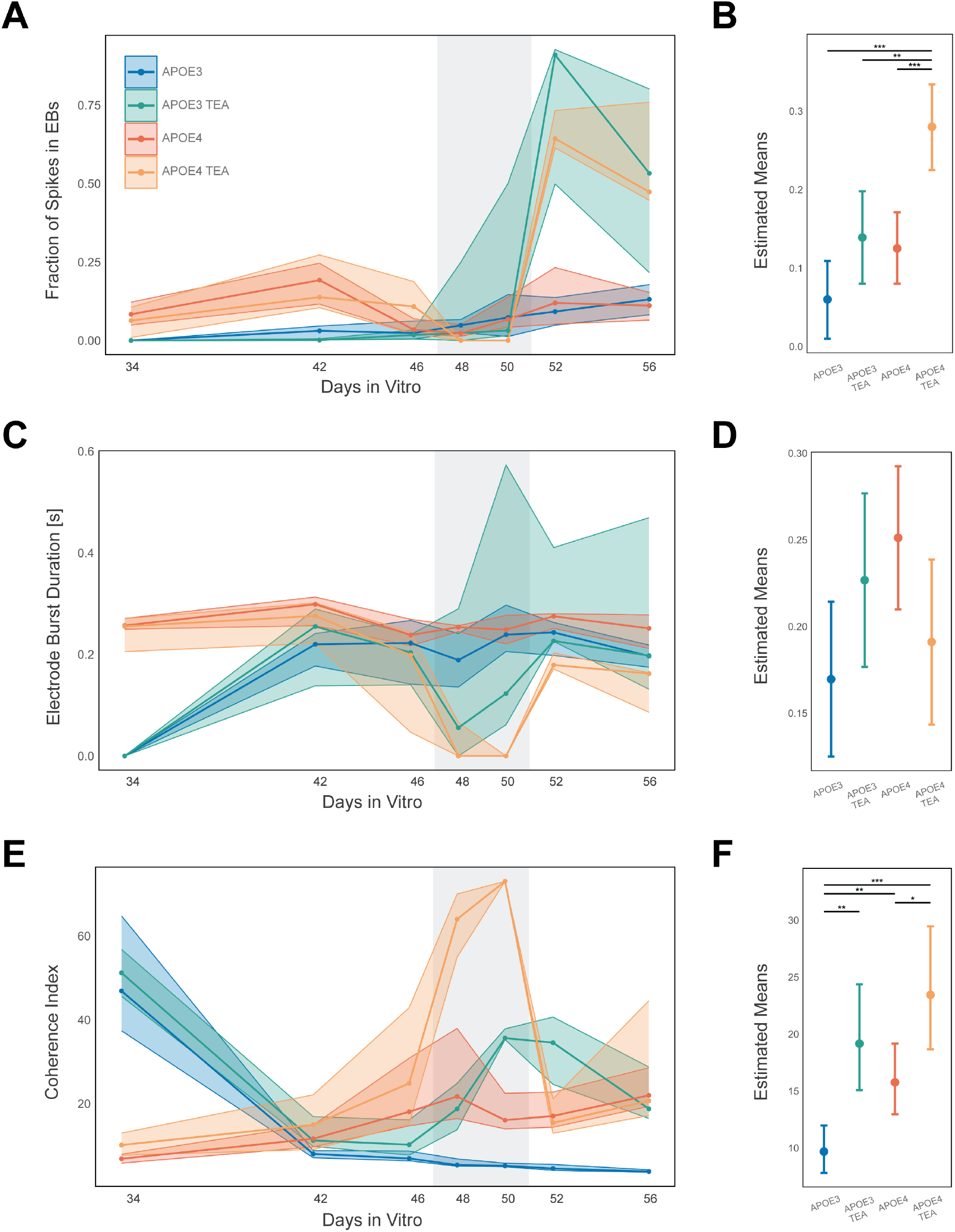
Network activity in ion-modulated (tetraethylammonium; TEA) and untreated control APOE3 and APOE4 networks. (**A-B**) Fraction of spikes in electrode bursts (EBs), (**C-D**) Electrode burst duration in s, and (**E-F**) Coherence index measuring network synchronization. Left panel: Neuronal network activity over time. Solid lines and circles illustrate median value for all networks in each group. Coloured shaded areas show upper and lower quartile. Vertical shaded gray areas indicate time points of treatment (47, 49, and 51 days in vitro). Group legend is shared between figures. Right panel: Generalized linear mixed models of estimated means with whiskers showing 95% confidence interval for each electrophysiology metrics. See Supplementary Table S3 for n per condition and time point. ***p<.001, **p<01, *p<.05

On the final recording before ion modulation, all groups showed approximately equal electrode burst durations (Fig. 6C-D). Ion modulation temporarily reduced the burst duration in both APOE3 and APOE4 networks. One day following the final perturbation, burst duration in both groups increased towards the levels observed in untreated networks, and this effect was stable until the subsequent final recording at 56 DIV. The longitudinal burst duration slopes did not significantly differ between any of the groups (Supplementary Table S2).

Given the marked reduction in the number of active electrodes following ion modulation (Fig. 5E-F), the coherence index was used to assess network synchronization (Fig. 6E-F) (Wang, 2002). While the coherence index showed a time-dependent reduction in APOE3 control networks, it slightly increased over time in APOE4 control networks (p<.01). Ion modulation increased the coherence index of both groups during perturbation, with the strongest effect observed in APOE4 networks. This effect was transient, and one and five days after the final TEA application, the coherence index of ion-modulated APOE4 networks was comparable to that of the untreated APOE4 control networks. For APOE3 networks, the increased coherence index during perturbation was still evident one day following the final TEA application, however, five days later, at the final recording at 56 DIV, the coherence index of ion-modulated APOE3 networks was comparable to that of untreated APOE3 networks.

## Discussion

Compared with isogenic APOE3 controls, APOE4 networks exhibited progressive hypoactivity, reduced synaptic AMPAR levels, and increased network synchronization. A time-dependent shift from elevated to reduced firing rate and number of active electrodes in APOE4 coincided with the emergence of excessive structural clustering and reduced neurite branching, highlighting the tight coupling between structural and functional deterioration in neurodegenerative disease. Synaptic engagement by ion modulation partially restored APOE4 network function by stabilizing AMPAR levels and producing lasting increases in spike amplitude and firing rate, demonstrating that APOE4 networks retain some capacity for functional reorganization despite their underlying vulnerabilities. However, the elevated network synchronization and aberrant network morphology of APOE4 were not restored by ion modulation, indicating that recovery of local neuronal function did not extend to global network organization.

APOE4 networks showed a time-dependent reduction in firing rate and the number of active electrodes compared with isogenic APOE3 controls (Fig. 5). Together with our findings of reduced synaptic AMPAR expression in APOE4 networks at 52 DIV (Fig. 4), these results indicate impaired excitatory synaptic transmission, to which AMPARs are major contributors (Caya-Bissonnette and Béïque, 2024). Consistent with this, we and others have previously demonstrated altered excitatory-inhibitory balance in human APOE4 networks, accompanied by dysregulation of synaptic organization and signalling, including glutamatergic synapses (Groenlie et al., 2026) (Algamal et al., 2022) (Burns et al., 2026). Importantly, however, disrupted network activity in AD is highly dynamic and depends on both disease stage and network state, with excitatory and inhibitory drive changing over time as compensatory mechanisms emerge (Burns et al., 2026) (Gabitto et al., 2024). This dynamic reorganization may also explain the apparently conflicting reports of hyper- and hypoexcitability in the AD literature.(Ranasinghe et al., 2025) (Quiroz et al., 2010) (Bookheimer et al., 2000) (Castanho et al., 2025) (Ereira et al., 2024) (Petrella et al., 2011) (Lin et al., 2018). The dynamic nature of network alterations in AD is reflected in our data by a time-dependent shift in network activity from 34 to 42 DIV. During this period, APOE4 networks switched from having higher firing rates and a higher number of active electrodes than APOE3 networks, to a hypoactive state characterized by reduced firing rates and fewer active electrodes at 42 DIV. Together with our previous findings of comparable time-dependent shifts in network dynamics and excitatory-inhibitory balance in AD (Groenlie et al., 2026) but also ALS networks (Kollstrøm et al., 2025a), these findings demonstrate the value of longitudinal electrophysiological investigations of human neuronal networks in vitro for identifying early transitions in network function that are difficult to detect in patients.

In addition to the reduced firing rate and number of active electrodes in APOE4 networks, we identified an increased coherence index compared with isogenic APOE3 networks, indicating elevated network synchronization. This is consistent with previous findings of excessive synchrony in cellular models of both AD (Weir et al., 2024) and ALS (Kollstrøm et al., 2025b). We have previously reported reduced redundancy and increased assortativity in APOE4 networks, suggesting a greater reliance on a small number of critical pathways for information transfer (Groenlie et al., 2026). Within this context, the progressive reduction in the number of active electrodes and increased synchronization observed in APOE4 networks in the present study may reflect an homeostatic reconfiguration of the network dynamics towards a state in which the remaining neuronal activity becomes increasingly synchronized, thereby preserving information transfer despite declining local activity (Chowdhury and Hell, 2018) (Turrigiano, 2008) (Weir et al., 2024). Similarly, the increased fraction of spikes in bursts in APOE4 networks may further support information transfer despite the overall low activity (Lisman, 1997) (Csicsvari et al., 1998) (Kobayashi et al., 2024). This compensatory homeostatic scaling, however, characterized by fewer active electrodes but high global synchronization, may increase susceptibility to the spread of AD-pathology (Liu et al., 2012) (Wang et al., 2017), demonstrating that progressive functional reorganization and compensatory synchronization contribute to the emergence of a vulnerable network state in APOE4 networks.

One day before the identified shift in firing rate and number of active electrodes at 42 DIV, light microscopy revealed initial formation of excessively large clusters in APOE4 networks (Supplementary Fig. S1). These morphological changes were absent at the previous imaging time point at 27 DIV, suggesting that excessive clustering developed during the same time period as the shift in network activity. Aberrant growth and structural organization of neuronal networks strongly affect their functional network dynamics (Weir et al., 2024) (Ballesteros-Esteban et al., 2023). For instance, the extent of dendritic branching contributes to the integrative capacity and plasticity of neuronal networks (Desai-Chowdhry et al., 2023), and neurons with more extensive dendritic branches can achieve splay states for excitatory coupling, which neurons with fewer branches cannot (Gowers and Schreiber, 2024). In the context of neurodegenerative disease, Kollstrøm et al. (2025a) reported aberrant network morphology coinciding with hyperexcitability in ALS networks, and Weir et al. (2024) showed that neurite retraction and clustering correlated with reduced firing of tau-perturbed rat cortical neurons. Together, these findings suggest that the close link between structural and functional network defects in AD may explain the identified coinciding structural and functional deterioration of APOE4 networks in the present study.

Although the causal mechanism underlying the progressive impairments of APOE4 networks cannot be fully resolved from our data, several potential mechanisms are consistent with our observations. Impaired neurite outgrowth and repair has been demonstrated following APOE4 exposure in multiple model systems (Nathan et al., 1995) (Nathan et al., 2002) (Toro et al., 2021) (Hussain et al., 2013) (Nathan et al., 1994), and the structural and electrophysiological profiles of our APOE4 networks share notable similarities with those of networks perturbed with human tau (Weir et al., 2024). In the present work, AT8-positive tau was detected in the large APOE4 clusters, raising the possibility that tau pathology contributes to axonal retraction and clustering (Weir et al., 2024). We did not quantify Aβ or phosphorylated tau levels in the present study, as the asynchronous maturation typical of iPSC-derived networks makes individual cells occupy different developmental and pathological states at any given time point, introducing systematic bias into population-level comparisons. Apart from the expression of these classical pathological hallmarks of AD, impaired protein clearance by APOE4 astrocytes, a documented deficit (Wang et al., 2021) (Eisenbaum et al., 2024) (Fleeman and Proctor, 2021), could contribute to structural breakdown and impaired branching. Moreover, isoform-dependent differences in APOE-mediated lipid transport (Chen et al., 2025b) (Pinals and Tsai, 2022) may impair membrane homeostasis and ion channel function (Carlo, 2013), thereby disrupting cell integrity and adhesion (Groenlie et al., 2026), altering electrochemical signalling, and contributing to excessive water influx and cell swelling (Rungta et al., 2015) (Steffensen et al., 2015) (Lemale et al., 2022). Together, these findings support a strong relationship between structural and functional deterioration and declining network function in APOE4 networks, while suggesting that multiple interacting mechanisms likely converge to drive the observed network reorganization.

Ion modulation reduced dendritic branching and increased the average neurite length in APOE3 networks (Fig. 3). This aligns well with our previous findings following TEA-treatment of human ALS neurons, in which ion modulation similarly reduced neurite branching and increased neurite length (Kollstrøm et al., 2025b). Interestingly, Kollstrøm et al. (2025b) also identified proteomic upregulation of pathways involved in actin and cytoskeletal organization following ion modulation by TEA. This observation may help explain why a similar effect was not observed in our APOE4 networks, as we have previously identified dysregulation of proteins involved in the actin cytoskeleton, actin binding, and dendritic tree-related proteins in human APOE4 neurons (Groenlie et al., 2026), supporting previous reports that the networks’ capacity for adaptive structural reorganization is impaired (Toro et al., 2021). The excessive clustering observed in APOE4 networks may have further limited their structural plasticity. Future studies applying ion modulation before the emergence of this aberrant morphological phenotype may help determine whether early intervention may prevent the structural abnormalities observed in APOE4 networks.

Although ion modulation had different effects on the structural characteristics of APOE3 and APOE4 networks, functional effects were observed in both groups, including increased spike amplitude following perturbation, consistent with TEA’s mechanism of action. TEA prolongs the action potential duration by blocking potassium channels required for potassium efflux and action potential repolarization (Aniksztejn and Ben-Ari, 1991b) (Huang and Malenka, 1993) (Huber et al., 1995) (Aniksztejn and Ben-Ari, 1991a) (Paulsen et al., 1990). By prolonging membrane depolarization, TEA enhances presynaptic calcium influx and neurotransmitter release, consistent with the increased spike amplitude observed following ion modulation. Despite temporary increases in spike amplitude and firing rates after the perturbation period, the number of active electrodes was strongly and durably reduced in both groups during and after ion modulation. In other words, potassium channel blockage reduced the number of active synapses while increasing the activity of the remaining active synapses, consistent with synaptic scaling (Lu et al., 2025) (Turrigiano, 2008). This pattern is consistent with a homeostatic response in which the remaining active synapses compensate for reduced network participation. In APOE4 networks, this resulted in elevated firing rate one day after the final perturbation, together with stable GluR1 levels despite the reduced number of active electrodes. Since AMPAR levels were quantified across the entire network, an increase in overall AMPAR levels would not be expected given the substantial reduction in the number of active electrodes. Instead, the preservation of AMPAR levels despite a reduced number of active electrodes, together with increased firing rate, is consistent with compensatory upscaling of the remaining active synapses following TEA-induced silencing of parts of the network. These results demonstrate that despite underlying structural and functional vulnerability, APOE4 networks retain a capacity for adaptive synaptic plasticity.

Ion-modulated APOE3 networks similarly exhibited a sustained reduction in the number of active electrodes, yet a corresponding increase in firing rate did not emerge until 56 DIV. The suppressed firing rate observed at 52 DIV is consistent with the reduced AMPAR levels detected in TEA-treated APOE3 networks at this time point, leaving open the possibility that synaptic scaling operates by the same mechanism in APOE3 as in APOE4 networks, albeit with a delayed onset. This temporal divergence in the spiking dynamics between the two groups is further reflected in the spike amplitude, which in APOE4 networks remained persistently elevated following ion modulation, whereas in APOE3 networks it peaked at 52 DIV before returning to levels comparable to untreated controls by 56 DIV as the drug distribution or compensatory conductances reassert control, consistent with previous studies reporting dose-dependent and reversible effects of TEA (Scheller et al., 1998) (Barrett et al., 1988). This temporal divergence may reflect differences in the underlying network state at the time of the perturbation together with isoform-specific metabolic capacity, which could influence the timing and magnitude of the electrophysiological response to ion modulation.

Network synchrony, as measured by the coherence index, tells a complementary story. APOE3 networks exhibited a moderate but sustained increase in response to TEA, whereas APOE4 displayed a pronounced but transient surge confined largely to the perturbation phase. This divergence may reflect an unbalanced overcorrection in APOE4, where the silencing of already low-firing synapses elicits a disproportionate compensatory synchronization response that may transiently preserve network function (Chowdhury and Hell, 2018) (Turrigiano, 2008) (Weir et al., 2024). The transient synchronization peak in APOE4, combined with reduced neurite branching and aberrant network morphology, suggests that global information transfer is fundamentally compromised - with fewer neurite branches to propagate synchronous activity across the network, efficient global network coordination cannot be sustained (Desai-Chowdhry et al., 2023) (Gowers and Schreiber, 2024) (Groenlie et al., 2026). Impaired glial support from APOE4 astrocytes (Chen et al., 2025b) (Zhang et al., 2025) may further contribute to this, as astrocytes are strongly involved in the integration and synchronization of neuronal activity (Deemyad et al., 2018) (Lendemeijer et al., 2024) (Kumar et al., 2020) (Kanakov et al., 2019). In contrast, the lasting increases in spike amplitude and firing rate in APOE4 networks following ion modulation indicate that, at the level of individual spikes, the capacity for synaptic plasticity is at least partially preserved despite persistent impairment of global network coordination.

## Conclusion

We demonstrate that APOE4 networks exhibit emerging structural deterioration characterized by excessive clustering and reduced neurite branching. Simultaneously, APOE4 networks undergo a time-dependent transition from hyperactivity to hypoactivity, accompanied by reduced synaptic AMPAR expression and excessive synchronization, the latter consistent with homeostatic reconfiguration in response to reduced local spiking activity, albeit at the cost of increased vulnerability to pathological spread. Synaptic engagement by ion modulation partially restored the local firing dynamics and stabilized AMPAR levels, indicating that APOE4 networks retain some capacity for functional adaptation despite underlying structural and functional vulnerability. Yet, the persistence of unstable synchronization and aberrant network morphology following ion modulation revealed that recovery of local neuronal function was not accompanied by restoration of the global network organization. Together, these findings show that APOE4 networks retain a capacity for adaptive plasticity enabling restoration of local neuronal function, but that this recovery does not extend to higher-order network organization.

## AUTHORS’ CONTRIBUTIONS

**MBG:** Conceptualisation, Methodology, Software, Formal Analysis, Investigation, Data Curation, Writing - Original Draft, Writing - Review & Editing, Visualisation. **AMK:** Methodology, Software, Formal Analysis, Investigation, Writing - Original Draft, Writing - Review & Editing, Visualisation. **AS:** Conceptualisation, Resources, Writing - Review & Editing, Supervision, Funding acquisition. **IS:** Conceptualisation, Resources, Writing - Review & Editing, Supervision, Funding acquisition.

## ACKNOWLEDGEMENTS

Western blot was performed by the Proteomics and Modomics Experimental Core (PROMEC), Norwegian University of Science and Technology (NTNU) and The Central Norway Regional Health Authority. This facility is a member of the National Network of Advanced Proteomics Infrastructure (NAPI), which is funded by the Research Council of Norway INFRASTRUKTUR-program (project number: 295910). Data storage and handling is supported under the NIRD/Notur project NN9036K.

## ETHICS APPROVAL AND CONSENT TO PARTICIPATE

Ethical concerns regarding cell donor privacy do not arise in the present work, as all iPSC lines were obtained already anonymised.

## CONSENT FOR PUBLICATION

Not applicable.

## AVAILABILITY OF DATA AND MATERIALS

The datasets used and/or analysed during the current study are available from the corresponding author on reasonable request.

## LIST OF ABBREVIATIONS

AD: Alzheimer’s disease
ALS: Amyotrophic lateral sclerosis
AMPAR: α-amino-3-hydroxy-5-methyl-4-isoxazolepropionic acid receptor
APOE: Apolipoprotein E **A**β: Amyloid-β
DIV: Days in vitro
iPSCs: Induced pluripotent stem cells
LTP: Long-term potentiation
MEA: Microelectrode array
NMDAR: N-methyl-D-aspartate receptor
NTNU: Norwegian University of Science and technology
PROMEC: Proteomics and Modomics Experimental Core Facility
TEA: Tetraethylammonium

