## Supplementary Materials for "Ion modulation restores local synaptic function in APOE4 neuronal networks despite persistent impairments in global network organization"

**This file contains:**

Figure S1
Tables S1-S3

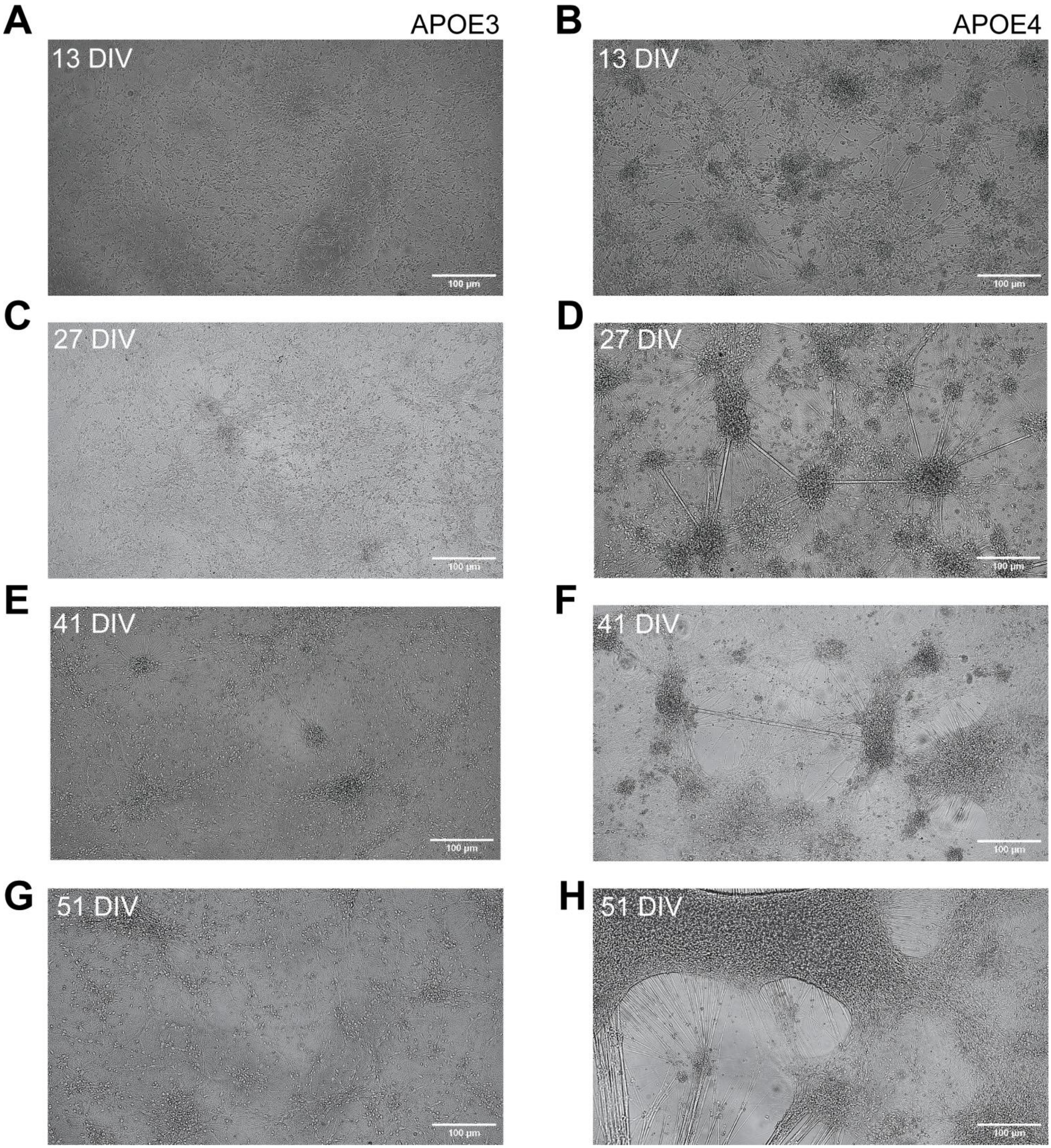


**Figure S1.** Representative light microscopy images of developing APOE3 and APOE4 neuronal networks. Left panel displays APOE3 networks. Right panel displays APOE4 networks. (**A-B**) 13 days in vitro (DIV). (**C-D**) 27 DIV. (**E-F**) 41 DIV. (**G-H**) 51 DIV. Scale bar 100 um.

**Table S1.** Estimated Means and Selected Target Distributions and Link Functions for the Electrophysiology Metrics.

| Measure | Group | Estimated means | SE | Lower 95% CI | Upper 95% CI | Distribution (link function) |
|---|---|---|---|---|---|---|
| Amplitude (mV) | APOE3 | .015 | .001 | .014 | .017 | Normal (identity) |
| Amplitude (mV) | APOE3 TEA | .018 | .001 | .016 | .020 | |
| Amplitude (mV) | APOE4 | .017 | .001 | .016 | .019 | |
| Amplitude (mV) | APOE4 TEA | .018 | .001 | .016 | .020 | |
| Normalized firing rate (Hz) | APOE3 | .231 | .028 | .181 | .294 | Gamma (log) |
| Normalized firing rate (Hz) | APOE3 TEA | .144 | .020 | .109 | .190 | |
| Normalized firing rate (Hz) | APOE4 | .092 | .010 | .073 | .115 | |
| Normalized firing rate (Hz) | APOE4 TEA | .117 | .015 | .090 | .152 | |
| Active electrodes | APOE3 | 38.667 | 1.663 | 35.333 | 42.000 | Normal (identity) |
| Active electrodes | APOE3 TEA | 24.320 | 1.895 | 20.539 | 28.101 | |
| Active electrodes | APOE4 | 32.289 | 1.536 | 29.200 | 35.378 | |
| Active electrodes | APOE4 TEA | 25.576 | 1.800 | 21.981 | 29.172 | |
| Fraction spikes in EBs | APOE3 | .060 | .025 | .010 | .109 | Normal (identity) |
| Fraction spikes in EBs | APOE3 TEA | .139 | .030 | .080 | .198 | |
| Fraction spikes in EBs | APOE4 | .125 | .022 | .080 | .171 | |
| Fraction spikes in EBs | APOE4 TEA | .280 | .027 | .225 | .334 | |
| EB duration | APOE3 | .170 | .022 | .125 | .214 | Normal (identity) |
| EB duration | APOE3 TEA | .227 | .025 | .177 | .277 | |
| EB duration | APOE4 | .251 | .020 | .210 | .293 | |
| EB duration | APOE4 TEA | .191 | .024 | .143 | .239 | |
| Coherence index | APOE3 | 9.678 | 1.031 | 7.817 | 11.982 | Gamma (log) |
| Coherence index | APOE3 TEA | 19.163 | 2.308 | 15.069 | 24.369 | |
| Coherence index | APOE4 | 15.762 | 1.533 | 12.961 | 19.168 | |
| Coherence index | APOE4 TEA | 23.445 | 2.680 | 18.656 | 29.463 | |

EB, electrode burst; SE, standard error; CI, confidence interval

**Table S2.** Pairwise Comparisons for the Electrophysiology Metrics.

| Measure | Comparison | Estimate | SE | t | df | p-value[α] |
|---|---|---|---|---|---|---|
| Amplitude (mV) | APOE3 TEA - APOE3 | .003 | .001 | 2.221 | 50.720 | .154 |
| Amplitude (mV) | APOE4 - APOE3 | .002 | .001 | 1.972 | 38.357 | .224 |
| Amplitude (mV) | APOE4 - APOE3 TEA | -.001 | .001 | -.537 | 49.207 | 1.000 |
| Amplitude (mV) | APOE4 - APOE4 TEA | -.001 | .001 | -.594 | 48.380 | 1.000 |
| Amplitude (mV) | APOE4 TEA - APOE3 | .003 | .001 | 2.304 | 49.980 | .152 |
| Amplitude (mV) | APOE4 TEA - APOE3 TEA | .000 | .001 | .041 | 59.368 | 1.000 |
| Normalized firing rate (Hz) | APOE3 TEA - APOE3 | -.086 | .034 | -2.516 | 54.787 | .059 |
| Normalized firing rate (Hz) | APOE4 - APOE3 | -.139 | .030 | -4.658 | 49.602 | .000 |
| Normalized firing rate (Hz) | APOE4 - APOE3 TEA | -.052 | .022 | -2.324 | 59.969 | .071 |
| Normalized firing rate (Hz) | APOE4 - APOE4 TEA | -.025 | .019 | -1.361 | 54.910 | .358 |
| Normalized firing rate (Hz) | APOE4 TEA - APOE3 | -.113 | .032 | -3.557 | 52.467 | .004 |
| Normalized firing rate (Hz) | APOE4 TEA - APOE3 TEA | -.027 | .025 | -1.072 | 63.297 | .358 |
| Active electrodes | APOE3 TEA - APOE3 | -14.347 | 2.522 | -5.690 | 62.052 | .000 |
| Active electrodes | APOE4 - APOE3 | -6.378 | 2.264 | -2.817 | 51.424 | .019 |
| Active electrodes | APOE4 - APOE3 TEA | 7.969 | 2.440 | 3.266 | 59.702 | .007 |
| Active electrodes | APOE4 - APOE4 TEA | 6.712 | 2.366 | 2.837 | 56.727 | .019 |
| Active electrodes | APOE4 TEA - APOE3 | -13.090 | 2.450 | -5.342 | 59.269 | .000 |
| Active electrodes | APOE4 TEA - APOE3 TEA | 1.257 | 2.614 | .481 | 66.595 | .632 |
| Fraction spikes in EBs | APOE3 TEA - APOE3 | .079 | .039 | 2.055 | 62 | .132 |
| Fraction spikes in EBs | APOE4 - APOE3 | .066 | .033 | 1.975 | 46 | .132 |
| Fraction spikes in EBs | APOE4 - APOE3 TEA | -.014 | .037 | -.366 | 60 | .716 |
| Fraction spikes in EBs | APOE4 - APOE4 TEA | -.154 | .035 | -4.353 | 55 | .000 |
| Fraction spikes in EBs | APOE4 TEA - APOE3 | .220 | .037 | 5.967 | 57 | .000 |
| Fraction spikes in EBs | APOE4 TEA - APOE3 TEA | .141 | .040 | 3.474 | 70 | .004 |
| EB duration | APOE3 TEA - APOE3 | .057 | .033 | 1.728 | 35.909 | .371 |
| EB duration | APOE4 - APOE3 | .082 | .030 | 2.746 | 28.735 | .062 |
| EB duration | APOE4 - APOE3 TEA | .024 | .032 | .766 | 34.208 | .909 |
| EB duration | APOE4 - APOE4 TEA | .060 | .031 | 1.943 | 32.078 | .304 |
| EB duration | APOE4 TEA - APOE3 | .021 | .032 | .669 | 33.888 | .909 |
| EB duration | APOE4 TEA - APOE3 TEA | -.036 | .034 | -1.044 | 39.011 | .909 |
| Coherence index | APOE3 TEA - APOE3 | 9.485 | 2.527 | 3.753 | 64.770 | .002 |
| Coherence index | APOE4 - APOE3 | 6.084 | 1.847 | 3.294 | 48.666 | .007 |
| Coherence index | APOE4 - APOE3 TEA | -3.401 | 2.770 | -1.228 | 60.055 | .449 |
| Coherence index | APOE4 - APOE4 TEA | -7.683 | 3.088 | -2.488 | 58.201 | .047 |
| Coherence index | APOE4 TEA - APOE3 | 13.767 | 2.872 | 4.794 | 61.372 | .000 |
| Coherence index | APOE4 TEA - APOE3 TEA | 4.282 | 3.537 | 1.211 | 64.661 | .449 |

EB, electrode burst; SE, standard error; df, degrees of freedom.

[α] Bonferroni adjusted

**Table S3.** Number of networks on microelectrode arrays included in the electrophysiology analysis.

| **Days in vitro** | ***n* APOE3** | ***n* APOE3 TEA** | ***n* APOE4** | ***n* APOE4 TEA** | **Total *N*** |
|---|---|---|---|---|---|
| 34 | 5 | 2 | 12 | 12 | 31 |
| 42 | 12 | 12 | 12 | 12 | 48 |
| 46 | 12 | 12 | 12 | 10 | 46 |
| 48 | 12 | 12 | 12 | 6 | 42 |
| 50 | 12 | 3 | 12 | 1 | 28 |
| 52 | 12 | 3 | 12 | 3 | 30 |
| 56 | 11 | 4 | 12 | 5 | 32 |
| **Total** | 76 | 48 | 84 | 49 | 257 |
